# Engineering the microstructure and spatial bioactivity of MAP scaffolds in vitro instructs neovascularization in vivo

**DOI:** 10.1101/2023.11.01.565182

**Authors:** Alexa R. Anderson, Eleanor L. P. Caston, Lindsay Riley, Long Nguyen, Dimitris Ntekoumes, Sharon Gerecht, Tatiana Segura

## Abstract

In tissues where the vasculature is either lacking or abnormal, biomaterials can be designed to promote vessel formation and enhance tissue repair. In this work, we independently tune the microstructure and bioactivity of microporous annealed particle (MAP) scaffolds to guide cell patterning in 3D and promote de novo assembly of endothelial progenitor-like cells into vessels. We implement both *in silico* characterization and *in vitro* experimentation to elucidate an optimal scaffold formulation for vessel formation. We determine that MAP scaffolds with pore volumes on the same order of magnitude as cells facilitate cell growth and vacuole formation. We achieve spatial control over cell spreading by incorporating adhesive microgels in well-mixed, heterogeneous MAP scaffolds. While we demonstrate that integrin engagement is the primary driver of network formation in these materials, introducing adhesive microgels loaded with heparin nanoparticles leads to the formation of vascular tubes after 3 days in culture. We then show *in vivo* that this unique scaffold formulation enhances vessel maturation in a wound healing model and instructs differential vascular patterning in the tumor microenvironment. Taken together, this work determines the optimal microstructure and ligand presentation within MAP scaffolds that lead to vascular constructs *in vitro* and facilitate neovascularization *in vivo*.

## 1. Introduction

It has been well established that the revascularization of damaged or ischemic tissue is essential for repair and regeneration. The body relies on two mechanisms for forming new vascular networks: angiogenesis (i.e., new vessel sprouting from pre-existing vessels) and vasculogenesis (i.e., de novo assembly of endothelial progenitor-like cells into vessels).^[1]^ Vasculogenesis is known for its role in establishing primitive vascular networks from angioblasts in early embryonic development; however, postnatal vasculogenesis also occurs with circulating progenitor cells in the contexts of tissue repair and tumor development.^[2]^ Vessel formation through mechanisms of both vasculogenesis and angiogenesis are highly regulated and are mediated by both biochemical and physical cues.

Endothelial cells (ECs) are the primary building block of blood vessels which makes them an essential component of *in vitro* models for vessel formation. In their simplest form, reductionist *in vitro* models of vessel formation include ECs in a 2D system. However, the extracellular matrix (ECM) and support cells (i.e., vascular smooth muscle cells, pericytes) are also instrumental players in vessel formation, maturity, and function.^[3]^ The ECM serves as the host for paracrine signaling molecules while also playing a pivotal role in blood vessel formation through cell-matrix interactions.^[4]^ Support cells provide an array of functions to assist in vessel formation and regulation, such as clearing the path for vessel growth, secreting growth suppression signals, and creating a physical barrier as they wrap around the vessel to improve barrier function.^[5]^ Increasing the complexity of *in vitro* models through 3D matrices and the addition of support cells have improved the longevity and function of the microvascular systems that can be formed *in vitro*.^[6]^

In tissues where the vasculature is either lacking or abnormal, biomaterial interventions which mimic the ECM can be designed to induce vessel formation and promote tissue repair.^[7]^ Traditionally, the field of tissue engineering has used materials loaded with biochemical signals, such as vascular endothelial growth factor-A (VEGF), to promote and guide vessel formation; however, porous scaffolds without biochemical signals have also been shown to promote vascularization in vivo.^[8]^ The porous architecture of biomaterials plays a key role in influencing cell infiltration and inducing vascularization by enabling the diffusion of nutrients and providing structural avenues for vessel ingrowth.^[9]^ Granular hydrogels are a class of biomaterial that inherently possess a tunable, porous architecture. These materials are composed of small, building block subunits that pack together to produce an interconnected, porous network. Microporous annealed particle (MAP) scaffolds are a unique subclass of granular hydrogels that incorporate an additional *in situ* crosslinking mechanism to interlink the hydrogel subunits.^[10]^ We and others have shown that MAP scaffolds support endogenous cell infiltration, and that this system is both modular and adaptable for different tissue-specific applications.^[10, 11]^ While the inclusion of growth factors in granular materials^[12]^ or extracellular vesicles in MAP scaffolds^[13]^ has been shown to enhance vascular infiltration in vivo, more ‘vanilla’ formulations of MAP scaffolds have also demonstrated improved vessel infiltration.^[14]^ However, the formation of mature vascular structures with lumens has yet to be reported from *in vitro* implementation of MAP scaffolds.

Thus far, *in vitro* investigation of the role of granular material properties on vessel formation have been focused on studying sprouting angiogenesis of endothelial cells (ECs) from multicellular spheroids in MAP scaffolds. These foundational works have demonstrated that incorporating heparin microgels (μgels) in polyethylene glycol (PEG) MAP scaffolds^[15]^ and including an interstitial hyaluronic acid (HA) matrix between HA μgels^[12]^ improves EC sprouting and migration into the scaffold. It has also been described that in granular materials, the ECs preferentially spread along the μgels but do not form lumen-like sprouts.^[12]^ These *in vitro* studies using multicellular spheroids have allowed us to understand the properties of granular hydrogels that can be manipulated to modulate angiogenic sprouting; however, the pro-vasculogenic properties of these materials have yet to be investigated.

Using the MAP scaffold platform, we were interested in determining which matrix properties have an instructive effect on the de novo assembly of endothelial progenitor-like cells into vessels (i.e., vasculogenesis) to inform *in vivo* material design. In particular, we aimed to use a reductionist model consisting of the ECs and our biomaterial matrix with the goal of producing vessel structures with lumens without requiring support cells. We hypothesized that both scaffold microarchitecture and bioactivity could be tuned to guide vascular morphogenesis. Lastly, we sought to investigate whether this approach which uses a simplified *in vitro* model of vasculogenesis would translate to an effect on vessel formation in the contexts of tissue repair and tumor development *in vivo*.

## 2. Results and Discussion

### 2.1. Pore size controls EC growth in MAP scaffolds

The base composition of the µgels used in this work was a hyaluronic acid (HA) backbone modified with norbornene (NB) functional handles. The µgels were internally crosslinked with an MMP-labile peptide sequence via thiol-ene click chemistry as shown in **Figure 1**, then covalently interlinked using an HA polymer modified with tetrazine (Tet) functional handles to form the MAP scaffold.^[16]^ To determine the effect of MAP scaffold microstructure on endothelial cell growth, we fabricated μgels of different sizes to alter the pore size between μgels. For these studies, we added 1 mM RGD peptide to the µgels via thiol-ene click chemistry, and we controlled the flow rates of our microfluidic droplet generator device to produce three populations of μgels: 50 μm, 80 μm, and 200 μm average diameter (**Figure 1B-E**). Due to the nature of granular materials, we expected the pore size between µgels to increase with increasing µgel size. To computationally assess the characteristics of granular materials formed by our HA µgels, we simulated particle scaffolds of rigid spheres that mimicked the size distributions of our µgels (**Figure S1A-D**). For these particle scaffolds, we implemented LOVAMAP software as described by Riley, et al.^[17]^ to characterize the porous microstructure. As the average μgel diameter increased from 50 μm to 80 μm to 200 μm, the median pore volume increased from 8.5 to 26.4 to 523 pL (**Figure 1F**) which demonstrated that we could produce different microstructures in MAP scaffolds by controlling µgel size.

**Figure 1.**
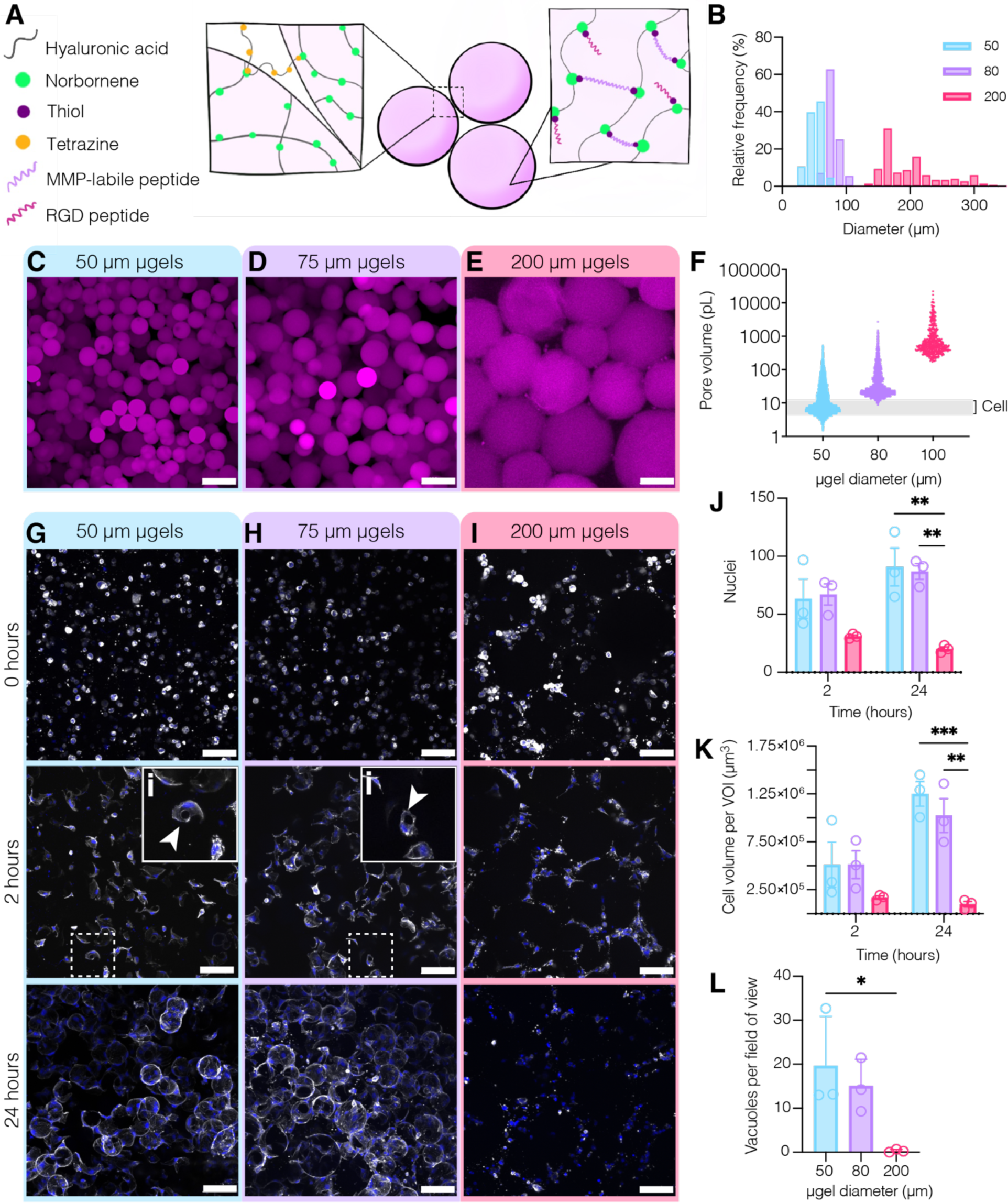
hECFC culture in MAP scaffolds with varying µgel size. (A) Schematic overview of the composition of MAP scaffolds used in this work. Three populations of µgels were produced with diameters of 50, 80, and 200 μm as shown by (B) the frequency distribution of µgel diameter and confocal images of MAP scaffolds with (C) 50 μm, (D) 80 μm, and (E) 200 μm µgels. (F) The pore volume for granular scaffolds composed of each µgel population was determined via computational analysis of rigid spheres with LOVAMAP. Representative maximum intensity projections (MIPs) spanning 100 μm of the 20x confocal Z-stack are shown for (G) 50 μm, (H) 80 μm, and (I) 200 μm over 24 h of culture (blue = DAPI, white = F-actin) (scale bar = 100 μm). Insets (i) highlight a cell with a vacuole as designated by the white arrow. (J) Cell counts across conditions over 24 h of culture. Two-way ANOVA with Tukey HSD was performed on *n* = 3 biological replicates with duplicate scaffolds. (K) Cell volume across conditions over 24 h of culture within a volume of interest. Two-way ANOVA with Tukey HSD was performed on *n* = 3 biological replicates with duplicate scaffolds. (L) Number of vacuoles per field of view across conditions after 2 h in culture. One-way ANOVA with Tukey HSD was performed on *n* = 3 biological replicates with duplicate scaffolds. Significance was reported at *p* < 0.05 (*), <0.01 (**), <0.005 (***), and <0.001 (****).

To study vasculogenesis in MAP scaffolds, we used human endothelial colony forming cells (hECFCs). hECFCs are adult endothelial progenitor cells which are highly proliferative and capable of forming vascular tubes *in vitro*.^[18]^ When cultured in MAP scaffolds, hECFCs spread along the µgels to form cell networks by 24 h (**Figure 1G-I**). We observed enhanced cell growth (via nuclei counts and cell volume) in the small and medium μgel conditions compared to large μgels (**Figure 1G-I**). More than this, the hECFCs formed vacuoles (an early indication of lumen formation) after only 2 hours of culture (**Figure 1E-G inset**), with 50 μm and 80 μm μgel MAP scaffolds boasting more vacuolating cells compared to 200 μm μgel scaffolds (**Figure 1J**). These findings demonstrate the role of biomaterial microarchitecture on EC response: in MAP scaffolds, smaller µgels (50-80 μm) promote early vacuole formation and ECFC growth as compared to large (200 μm) µgels. Interestingly, the pore volumes in MAP scaffolds composed of the smaller μgels shared the same order of magnitude as the volume of individual ECFCs at t=2 h (**Figure 1B, gray box**) as compared to the larger μgel scaffolds. As there were no significant differences between the 50 μm and 80 μm μgel groups, we used the medium μgel size (∼80 μm) for all subsequent studies (**Figure S2A-B**).

### 2.2. Distance between bioactive cues in heterogeneous MAP scaffolds is a function of both μgel proportion and size

In addition to structural cues, the presentation of adhesion ligands can be used to guide cell growth. We exploited the modularity of MAP scaffolds by mixing two µgel populations (Population A shown in pink and Population B shown in green) at different proportions to control the spatial distribution of bioactive cues within MAP scaffolds. This approach has been described previously for creating micro-islands of bioactive cues^[15, 19]^ as well as for supporting guest-host interlinking of MAP scaffolds^[20]^. We first validated that lyophilized µgels can be rehydrated and combined in varying proportions to yield scaffolds with consistent heterogeneous compositions. We created mixtures of 10% A + 90% B (**Figure 2A**), 25% A + 75% B (**Figure 2B**), 50% A + 50% B (**Figure 2C**), as well as 100% A (**Figure 2D**). While we were able to validate that the experimental proportions of each µgel population were consistent with the theoretical proportions for the scaffold (**Figure 2E-G**), this assessment did not capture the heterogeneity at higher resolution. As such, for a 50/50 mixture we also assessed the proportion of each µgel population within a subunit volume that would contain approximately four µgels (**Figure 2H**). While our early methods of mixing did not produce proportions centered at 50% (**Figure S2C**), we increased the duration of mixing to create heterogeneous MAP scaffolds with approximately 50% of each population within a subunit volume (**Figure 2I**), indicating that these scaffolds were well-mixed both globally and regionally within the material.

**Figure 2.**
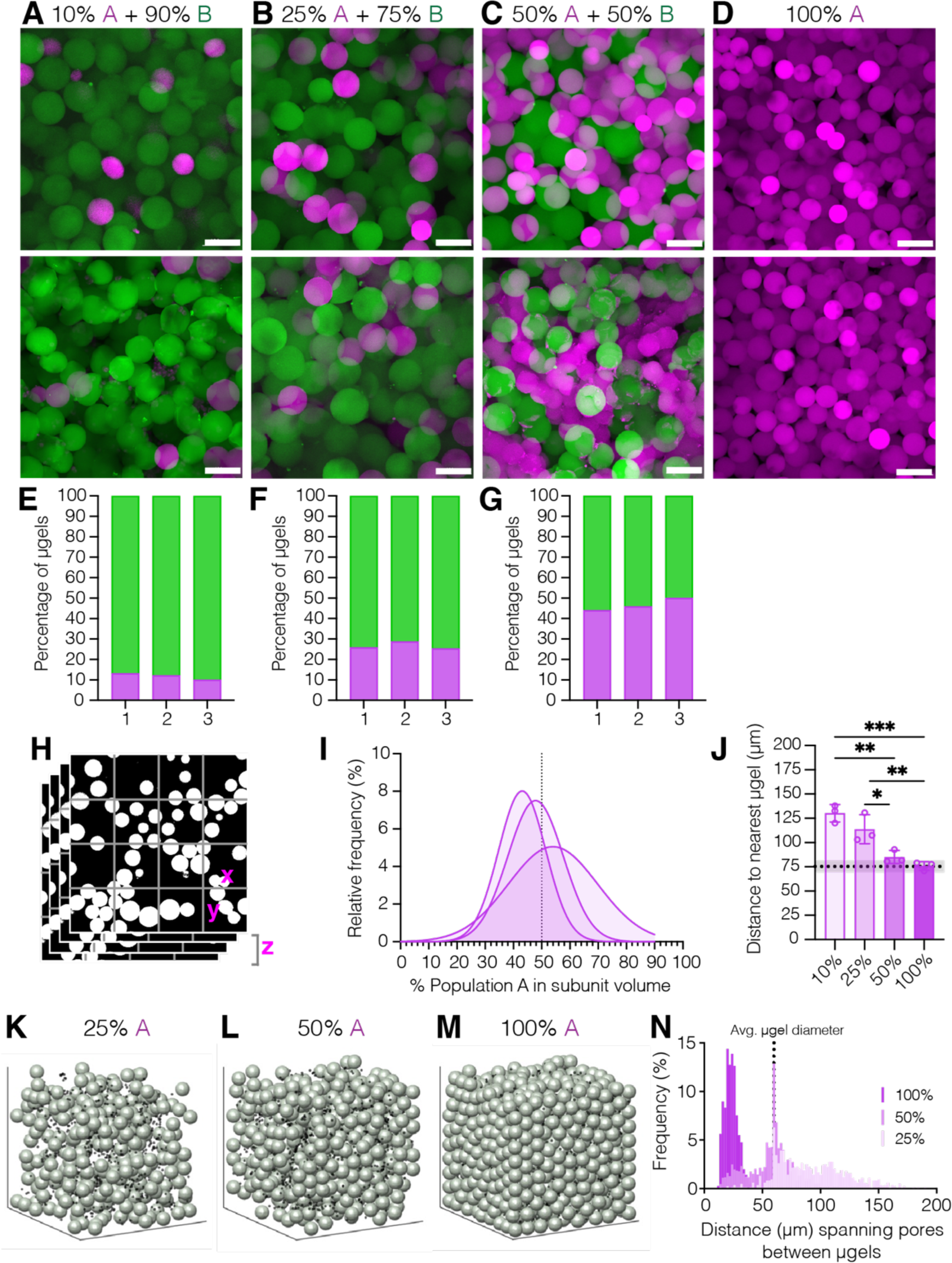
Heterogeneous mixtures of µgels in MAP scaffolds. Confocal images of MAP scaffolds comprised of (**A**) 10% A + 90% B, (**B**) 25% A + 75% B, (**C**) 50% A and 50% B, and (**D**) 100% A. The quantified percentage of µgels of each population for MAP scaffolds comprised of (**E**) 10% A + 90% B, (**F**) 25% A + 75% B, (**G**) 50% A and 50% B. (**H**) Confocal Z-stacks can be segmented into subunit volumes to quantify local mixing efficacy. (**I**) Frequency distribution of the percentage of Population A within a subunit volume of the confocal Z-stack of three independent MAP scaffolds composed of 50% A + 50% B. (**J**) Distance between µgels of the same species in heterogeneous mixtures. One-way ANOVA with Tukey HSD was performed on *n* = 3 biological replicates with duplicate scaffolds. Significance was reported at *p* < 0.05 (*), <0.01 (**), <0.005 (***), and <0.001 (****). (**K**) 25%, (**L**) 50%, and (**M**) 100% of Population A for simulated scaffolds of rigid spheres. Black dots indicate local center points within the empty space after Population B particles have been removed. (**N**) Frequency distribution of gap lengths between µgels in the simulated domains; annotating the average µgel diameter of Population B (dotted line).

With the goal of using μgel heterogeneity to control bioactivity, we hypothesized that the length scale between µgels of the same species would be important for cell engagement. To study the relationship between µgel heterogeneity and distance, we first quantified the average distance between µgel centroids using surface renders of microscope images. As expected, the distance between µgel centroids of the same species increased as the proportion of that population decreased (**Figure 2J**). In scaffolds with 100% of Population A, the average distance to the nearest neighbor was shown to be equivalent to the average μgel diameter, indicating that a μgel in this condition is touching its nearest neighbor of the same species.

We next used LOVAMAP to measure the size of gaps between µgel surfaces, rather than using µgel centroid coordinates. We simulated a particle scaffold of 60 µm diameter rigid spheres, then randomly removed particles representing Population B to analyze the gap lengths between µgels of Population A (**Figure 2K-M**). At 100% Population A, we saw that the distribution of gap lengths peaked at around 25 µm (**Figure 2N**). As heterogeneity was introduced, a new peak emerged at 60 µm, which maximized at 50% Population A. This finding illustrates that in well-mixed systems at 50% Population A, the average gap length equals the average size of Particle B. At 25% Population A, another subtle peak emerged at 120 µm, which reflects two Population B particles are touching. These results suggest that the length scale between heterogeneous µgels is a tunable metric that is a function of both µgel size and proportion.

### 2.3. Adhesive µgel populations drive EC patterning in MAP scaffolds

With the aim of guiding EC patterning, we hypothesized that cells would preferentially grow along adhesive µgels in a scaffold with a mixture of adhesive and non-adhesive µgels. Using LOVAMAP, we identified the shortest paths a cell network could take if it formed along adhesive µgels starting from the center of a MAP scaffold and moving radially to exit the scaffold (shown as thick lines in the 2D schematic in **Figure 3A**). These paths comprise a subset of the total paths that exit the scaffold. We performed repeated measures analysis across different proportions of adhesive versus plain µgels and found that the proportion of adhesive µgel paths reached a maximal plateau when there was approximately 50% or more adhesive µgels in the scaffold (**Figure 3F**). Based on previous reports of cells preferentially growing along µgel surfaces rather than forming tube-like structures, we hypothesized that including non-adhesive µgels would guide cells to assemble in the void space to bridge the adhesive µgels. We modified the LOVAMAP analysis to only consider paths of a cell network that consistently exposed cells to a heterogeneous mix of adhesive and plain µgels en route to exiting the scaffold (shown as thick lines in the 2D schematic in **Figure 3B**). We performed repeated measures across different proportions of adhesive versus plain µgels and found that the proportion of heterogeneous-µgel-paths relative to all possible exit-paths maximized when there was a 50/50 mixture of adhesive and non-adhesive µgels in the scaffold (**Figure 3J**).

**Figure 3.**
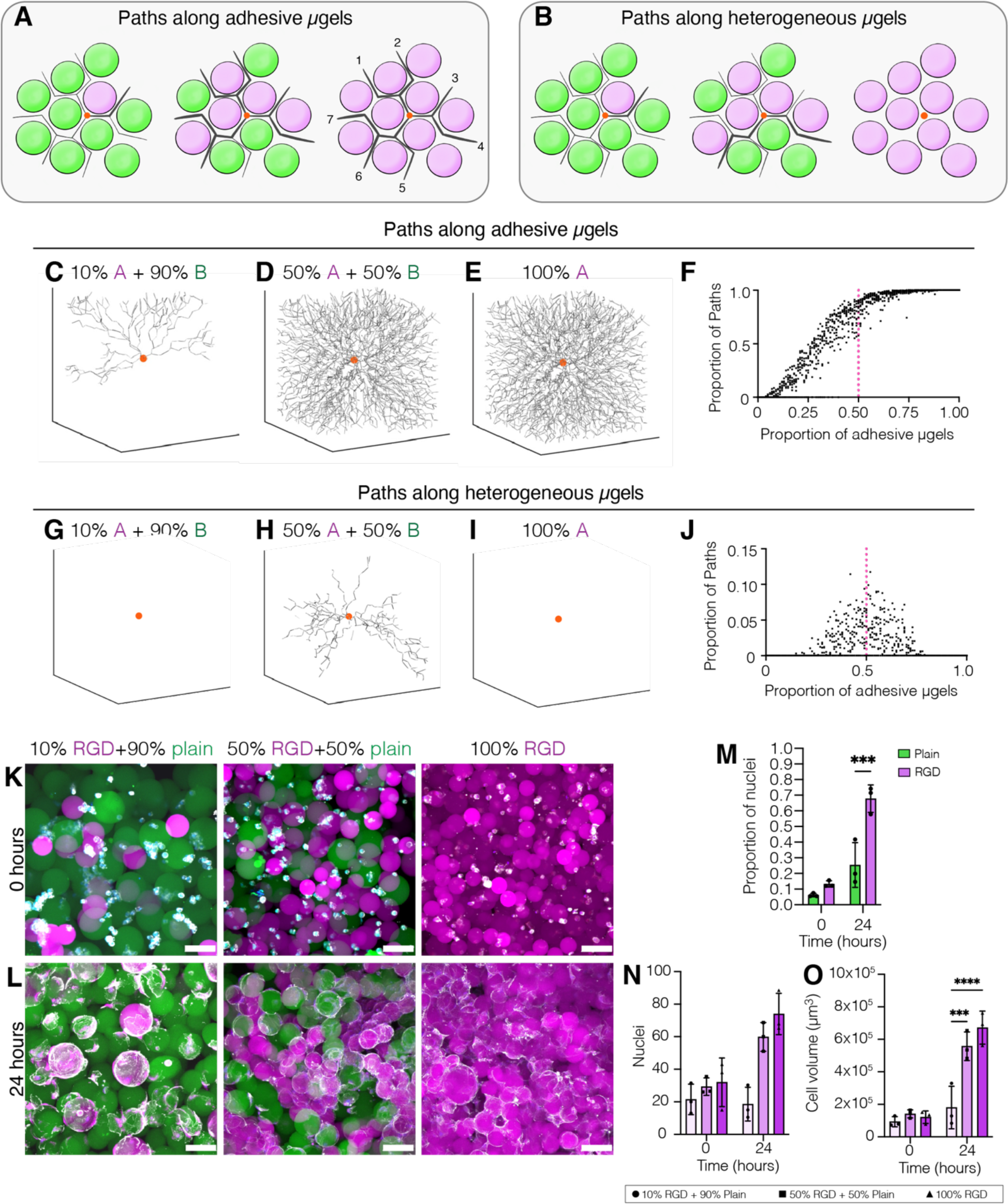
Directing cell growth using heterogeneous mixtures of µgels in MAP scaffolds. (**A**) 2D schematic of adhesive µgel paths identified by LOVAMAP analysis along which cell networks may form. In this simplified example, adhesive-µgel-paths (thick lines) must pass an adhesive µgel (magenta) at each step of the way. for cell networks forming along adhesive microgels (magenta). (**B**) 2D schematic of heterogeneous µgel paths identified by LOVAMAP analysis along which cell networks may form. In this simplified example, heterogeneous-µgel-paths (thick lines) must pass between an adhesive µgel (magenta) and a plain µgel (green) at each step of the way. Representative outputs of the network paths along adhesive µgels in heterogeneous MAP scaffolds composed of (**C**) 10% adhesive + 90% plain, (**D**) 50% adhesive + 50% plain, or (**E**) 100% adhesive µgels. (**F**) The proportion of paths identified for different heterogeneous combinations. Representative outputs of the network paths along combinations of adhesive and plain µgels in heterogeneous MAP scaffolds composed of (**G**) 10% adhesive + 90% plain, (**H**) 50% adhesive + 50% plain, or (**I**) 100% adhesive µgels. (**J**) The proportion of paths identified for different heterogeneous combinations. Confocal images of hECFCs in MAP scaffolds composed of 10% RGD + 90% plain, 50% RGD + 50% plain, and 100% RGD at (**K**) time = 0 and (**L**) time = 24 h. Scale bar = 100 µm. (**M**) Proportion of nuclei colocalized with either plain (green) or RGD (magenta) µgels over time. Increasing the proportion of adhesive µgels in the system resulted in increased cell growth, as quantified by (**N**) nuclei counts and (**O**) total hECFC volume per volume of interest. Two-way ANOVA with Tukey HSD was performed on *n* = 3 biological replicates with duplicate scaffolds. Significance was reported at *p* < 0.05 (*), <0.01 (**), <0.005 (***), and <0.001 (****).

To assess the response of hECFCs in heterogeneous MAP scaffolds, we combined different proportions of adhesive (1 mM RGD-modified) µgels (10%, 50%, 100%) versus plain µgels (90%, 50% 0%) and found that at t=0, cells were homogenously distributed in the scaffold (**Figure 3K**). However, over time cells localized with the µgels boasting the cell adhesion ligand (**Figure 3L-M**). This use of heterogeneous µgel populations allowed us to spatially guide cell growth in MAP scaffolds along the adhesive µgels. Increasing the proportion of RGD-modified µgels in the system resulted in increased cell growth, as quantified by nuclei counts (**Figure 3N**) and total hECFC volume (**Figure 3O**); however, there were not significant differences between the 50% RGD + 50% plain and 100% RGD scaffold conditions. These findings demonstrate that hECFC sprout formation over the first 24 hours in culture follows the LOVAMAP model of cell network growth along adhesive µgels.

### 2.4. Heparin nanoparticles drive EC patterning in homogeneously adhesive MAP scaffolds

In addition to using adhesivity, we hypothesized that incorporating a growth factor-binding moiety, such as heparin, in MAP would sequester growth factors in the local microenvironment to encourage EC spreading and increase the longevity of the cell networks. We produced µgels loaded with heparin nanoparticles (nH)^[21]^ to study this hypothesis, as it has been demonstrated that nanoparticles formulated from heparin i) lose their anticoagulation activity and ii) promote blood vessel formation when delivered in nanoporous HA hydrogels to the stroke infarct.^[22]^ We first combined different proportions of nH µgels (10%, 50%, 100%) versus plain µgels (90%, 50% 0%) and found that over time, hECFCs remained homogenously distributed in the scaffold (**Figure 4A-B**) and did not preferentially colocalize with either µgel population in the absence of adhesion ligand (**Figure 4C**). Additionally, cell growth did not increase over time without adhesion ligand in the system (**Figure 4D-E**).

**Figure 4.**
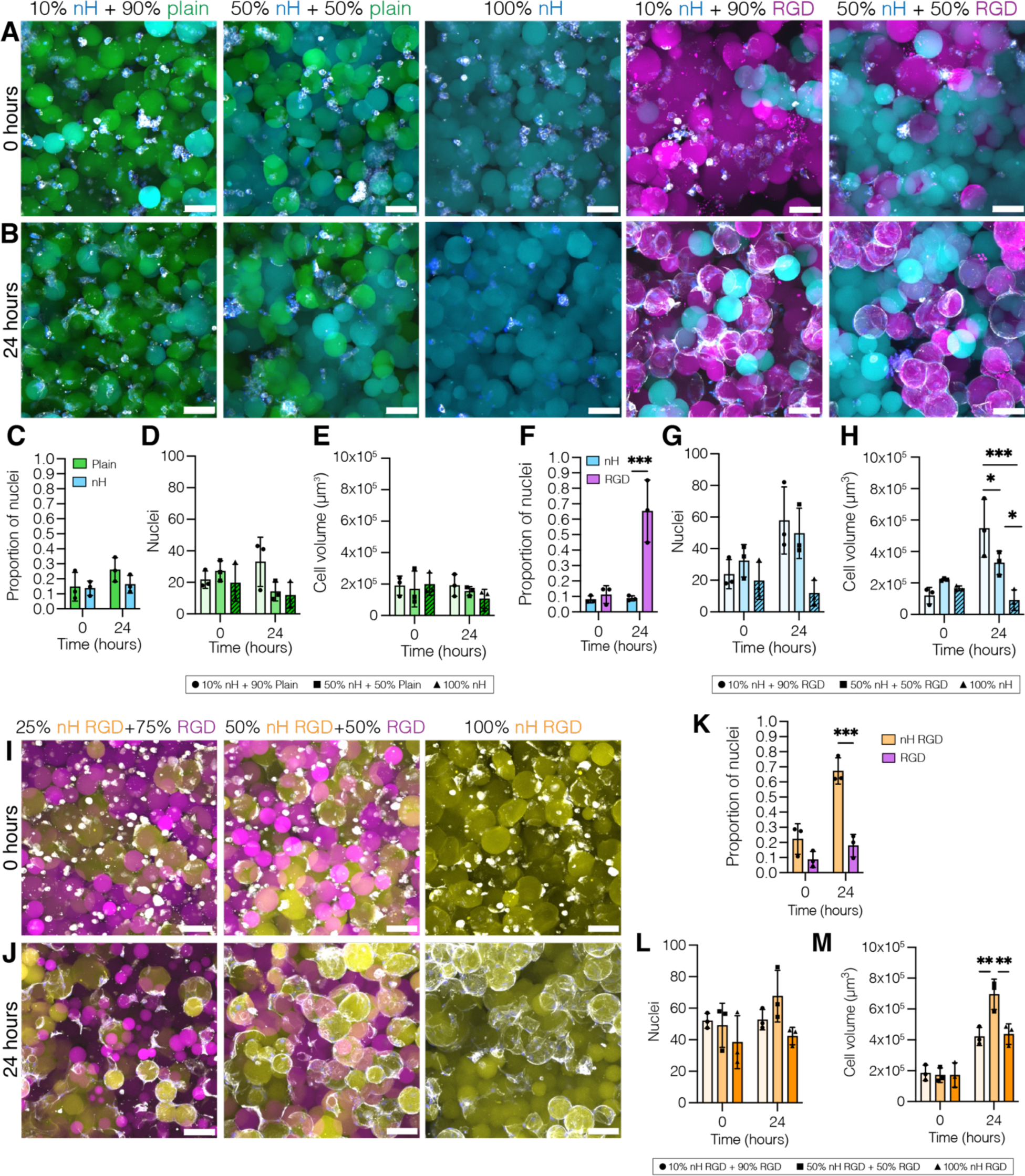
Confocal images of hECFCs in MAP scaffolds composed of 10% nH + 90% plain, 50% nH + 50% plain, 100% nH, 10% nH + 90% RGD, 50% nH + 50% RGD µgels at (**A**) time = 0 and (**B**) time = 24 h. Scale bar = 100 µm. (**C**) hECFCs did not preferentially localize with either population of µgels for the nH + plain conditions. Minimal changes were observed in (**D**) nuclei and (**E**) cell volume without RGD in the system. (**F**) Incorporating RGD in the system once again resulted in cells preferentially localizing with RGD-modified microgels after 24 hours in culture. Increasing RGD in the system resulted in increased cell growth, as quantified by (**G**) nuclei counts and (**H**) total hECFC volume per volume of interest. Confocal images of hECFCs in MAP scaffolds composed of 25% nH/RGD + 75% RGD, 50% nH/RGD + 50% RGD, and 100% nH/RGD µgels at (**I**) time = 0 and (**J**) time = 24 h. Scale bar = 100 µm. (**K**) Proportion of nuclei colocalized with either nH/RGD (orange) or RGD (magenta) µgels over time. Cell growth was quantified by (**L**) nuclei counts and (**M**) total hECFC volume per volume of interest. Two-way ANOVA with Tukey HSD was performed on *n* = 3 biological replicates with duplicate scaffolds. Significance was reported at *p* < 0.05 (*), <0.01 (**), <0.005 (***), and <0.001 (****).

We then reintroduced adhesive µgels into the system by combining different proportions of nH µgels (10%, 50%, 100%) with RGD-modified µgels (90%, 50% 0%) (**Figure 4A-B**). Similar to the results above for RGD vs plain µgels, cells are initially homogenously distributed in the scaffold but localize with the RGD-modified µgels over time (**Figure 4F**). Cell growth decreased as the proportion of adhesive µgels in the scaffold decreased (**Figure 4G-H**). These data show that over the first 24 hours, integrin binding facilitated by the RGD peptide was the primary driver of cell growth, and cell growth correlated with increasing proportion of RGD-modified µgels.

We thus decided to modify all µgels with RGD but change the proportion of µgels that contained nH. We combined heterogeneous populations of RGD-modified (75%, 50%, 0%) and nH/RGD-modified (25%, 50%, 100%) µgels for hECFC culture and found that at t=0, cells were homogenously distributed in the scaffold (**Figure 4I**). While cells did not localize with plain µgels containing nH (**Figure 4C, 4F**), the cells localized with the adhesive µgels containing nH after 24 hours (**Figure 4J-K**) which illustrates that nH can be used to direct cell growth in the presence of adhesive ligand. In terms of cell growth, we did not observe an increase in cell growth as we increased bioactivity; instead, the heterogeneous combination of 50% RGD and 50% nH/RGD µgels resulted in significantly more cell growth over 24 hours than the other combinations (**Figure 4M**).

### 2.5. Dual-functional µgels in MAP scaffolds support network maturation and lumen formation

While we observed comparable cell growth in all adhesive MAP scaffold conditions after 24 hours, we hypothesized that the incorporation of nH and its retention of growth factors would enhance network formation, complexity, and longevity in long-term cultures. We thus assessed hECFC networks after 3 days in culture for a range of heterogeneous MAP scaffold conditions with either RGD-modified µgels, nH µgels, nH/RGD-modified µgels, or some combination thereof (**Figure 5A-E**). In terms of nuclei counts (**Figure 5F**) and cell volume (**Figure 5G**), all combinations of RGD-modified + nH/RGD-modified µgels outperformed homogeneous RGD-modified MAP scaffolds as well as the heterogeneous MAP scaffolds with nH in plain µgels. hECFCs in 50% RGD and 50% nH/RGD µgels boasted an interconnected network on day 3 that is not observed in the other scaffold conditions. We quantified total network length (**Figure 5H**) and number of branches (**Figure 5I**) which reflected these observations and showed that the longest networks were from the 50% RGD and 50% nH/RGD µgel condition.

**Figure 5.**
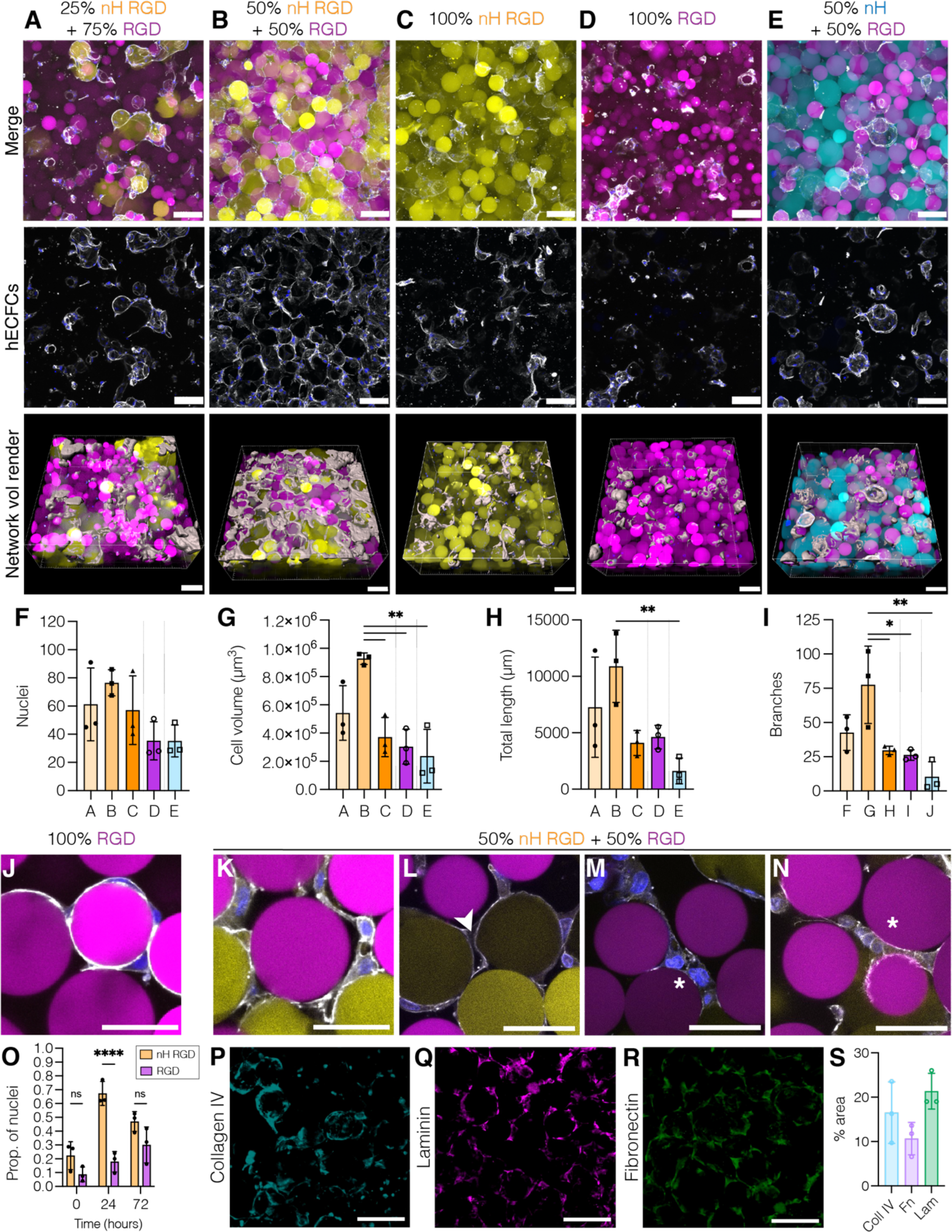
hECFC networks were assessed over 72 h in MAP scaffolds with (**A**) 25% nH/RGD + 75% RGD, (**B**) 50% nH/RGD + 50% RGD, (**C**) 100% nH/RGD, (**D**) 100% RGD, and (**E**) 50% nH + 50% RGD. Scale bar = 100 µm. Cell growth was quantified by (**F**) nuclei counts and (**G**) total hECFC volume per volume of interest. The (**H**) length and (**I**) branches of ECFC networks were quantified in 40 µm MIPs for each condition. One-way ANOVA with Tukey HSD was performed on *n* = 3 biological replicates with duplicate scaffolds. Significance was reported at *p* < 0.05 (*), <0.01 (**), <0.005 (***), and <0.001 (****). (**J**) hECFCs are shown wrapping around adhesive µgels in the 100% RGD condition. Scale bar = 50 µm. The mature markers in the hECFC networks cultured in MAP scaffolds with 50% RGD and 50% nH/RGD µgels are highlighted in single Z-slices from 40x images (**K-N**) and indicated by arrows for lumens and stars for cobblestone morphology. Scale bar = 50 µm. (**O**) Proportion of nuclei colocalized with either nH/RGD (orange) or RGD (magenta) µgels over time. Two-way ANOVA with Tukey HSD was performed on *n* = 3 biological replicates. Confocal images of hECFC networks in MAP scaffolds with 50% RGD + 50% nH/RGD µgels show deposition of ECM components (**P**) collagen IV, (**Q**) laminin, and (**R**) fibronectin in three separate scaffolds. (**S**) Quantification of the % area occupied by each ECM component quantified by fluorescence. One-way ANOVA with Tukey HSD was performed on *n* = 3 biological replicates. Significance was reported at *p* < 0.05 (*), <0.01 (**), <0.005 (***), and <0.001 (****).

As previously mentioned, 3D culture of ECs in MAP scaffolds was reported to result in cell spreading along μgel surfaces rather than forming tube-like structures.^[12]^ We similarly observed cells wrapping around μgels in the first 24 hours of culture, as well as after 72 hours of culture in adhesive MAP scaffolds (**Figure 5J**). However, we observed the formation of tube-like structures between microgels after 3 days in culture in heterogeneous MAP scaffolds with 50% RGD and 50% nH+RGD µgels (**Figure 5B**). In this scaffold composition, the tube structures formed by hECFCs exhibit markers of microvessel maturation. In particular, we observed cobblestone-like morphology of the hECFCs and interior lumens in the tube structures (**Figure 5K-N and Video #**). We showed above that after 24 hours, the hECFCs colocalized with the nH+RGD µgels; however, they did not preferentially colocalize with either population after 72 hours in culture (**Figure 5O**). These findings follow our second LOVAMAP model of cell network path predictions in which the optimal formulation for maximizing paths along heterogeneous μgels is 50% A + 50% B (**Figure 3B, H, J**).

ECs also deposit their own ECM (collagen IV, fibronectin, and laminin) as they assemble and form vascular structures^[23]^ which we hypothesized was occurring in the interstitial void space between the µgels in our VascMAP scaffolds. Through immunofluorescence staining, we detected the presence of collagen IV, fibronectin, and laminin deposited by hECFCs in the pores of MAP scaffolds with 50% RGD and 50% nH+RGD after 3 days in culture (**Figure 5P-S**).

### 2.6. Dual-functional µgels in MAP scaffolds support mature vessel formation in dermal wounds

As we have demonstrated the efficacy of this MAP formulation in supporting vessel formation *in vitro*, we sought to explore whether this response would translate *in vivo*. We performed a murine dermal wound healing model in which four full-thickness wounds were created on the back of each mouse with each wound receiving a different treatment (**Figure 6A**). In addition to testing our best-performing condition from our *in vitro* studies (HA µgels ∼80 µm in diameter with 50% RGD and 50% nH+RGD, termed VascMAP), we also included HA µgels ∼200 µm in diameter (Large HA) to assess the effect of microstructure, as well as PEG µgels ∼200 µm in diameter (Large PEG) to assess the effect of material on vessel response. We observed increased vessel formation (via CD31 staining) in all treatment groups compared to healthy skin at Day 7, but we observed significantly more vessels in the VascMAP condition compared to the untreated wound (**Figure 6B-E**, quantified in **Figure 6F**).

**Figure 6.**
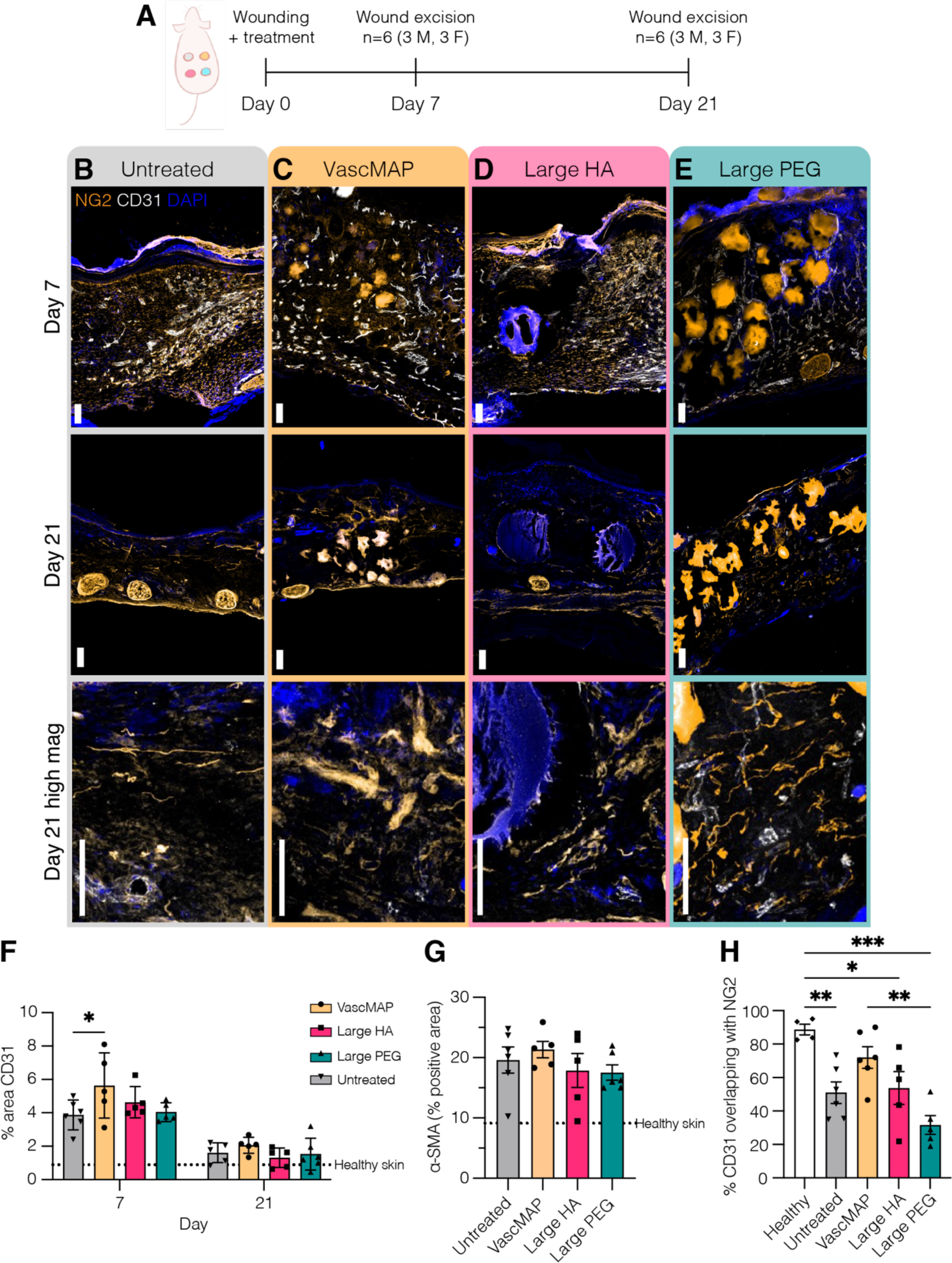
(**A**) Biomaterials introduced in a dermal wound healing model in SKH-1 mice were assessed at days 7 and 21 for vessel formation. Immunofluorescence staining of DAPI (blue), NG2 (orange), and CD31 (white) in cryosections from Days 7 and 21 for (**B**) untreated wounds or wound receiving (**C**) VascMAP, (**D**) Large HA MAP, or (**E**) Large PEG MAP. Scale bar = 100 µm. (**F**) The percent area positive for CD31 staining for each treatment group at days 7 and 21. Two-way ANOVA with Tukey HSD was performed to compare groups (*n* = 5-6). (**G**) The percent area positive for αSMA at day 21. One-way ANOVA with Tukey HSD was performed to compare groups. (**H**) The percent area positive of CD31 staining overlapping with positive NG2 staining for each treatment group at day 21. One-way ANOVA with Tukey HSD was performed to compare groups. Significance was reported at *p* < 0.05 (*), <0.01 (**), <0.005 (***), and <0.001 (****).

Sufficient perfusion of blood vessels is necessary for effective integration of biomaterials with the host tissue; however, ‘too much’ vessel formation can cause problems of its own. Vessel formation in the context of wound repair has been highly studied in skin wounds which shows a pattern of enhanced vessel growth followed by a period of decline as the wound resolves.^[24]^ While our goal is to design biomaterials that will become vascularized, the overabundance of blood vessels in wounds has been linked to fibrotic scar formation in skin and fibrosis in other tissue types as well.^[25]^ By Day 21, the heightened vessel formation resolved in all treatment groups to a level comparable to healthy skin (**Figure 6B-E**, quantified in **Figure 6F**). To determine if VascMAP was eliciting a more fibrotic response, we performed immunohistochemistry for α-smooth muscle actin (α-SMA, a marker for fibrogenic myofibroblasts) (**Figure S4H**). Staining for α-SMA at Day 21 showed all conditions elevated in comparison to healthy skin, but there were no significant differences in fibrogenesis amongst wounded groups (**Figure 6G**).

The rapid blood vessel formation that occurs during early wound healing leads to immature vessels with abnormal tortuosity and permeability; resultantly, the phase of vascular regression consists of pruning excess growth as well as the maturation of the remaining vessels.^[24]^ One indication of maturing vessel phenotype is the colocalization of mural cells with endothelial cells.^[26]^ At Day 21, we observed enhanced overlap of mural cells (via NG2 staining) with vessels in the VascMAP condition which was the only treatment that did not significantly differ from healthy skin in terms of mature vessels (**Figure 6G**). These findings demonstrate that VascMAP enhances vessel formation in the early stages of wound healing, and as the wound resolves and vessels regress, the remaining vessel structures exhibit more mature phenotypes than in the other biomaterial conditions.

### 2.7. Dual-functional µgels in MAP scaffolds alter vascular development in tumors

As previously mentioned, postnatal vasculogenesis occurs with circulating progenitor cells in the contexts of both tissue repair and tumor development.^[2]^ In comparison to tissues such as skin, tumors boast a high vascular density.^[27]^ Based on our findings in a dermal wound healing context, we sought to investigate whether VascMAP would have an effect in altering tumor vasculature in a murine model of glioblastoma, which is a type of highly vascularized solid tumor.^[28]^ We hypothesized that the heparin µgels in VascMAP would sequester endogenous vascular growth factors that are upregulated in the tumor microenvironment to instruct differential tumor vessel formation. For this model, we once again compared VascMAP to MAP comprised of large HA µgels and large PEG µgels. We inoculated CT2A mouse glioma cells in 5 µL of each biomaterial as well as a methylcellulose control in the mouse cortex (**Figure 7A**). We assessed tumor growth over 2 weeks via IVIS imaging and saw no significant differences in tumor growth amongst conditions (**Figure 7B**).

**Figure 7.**
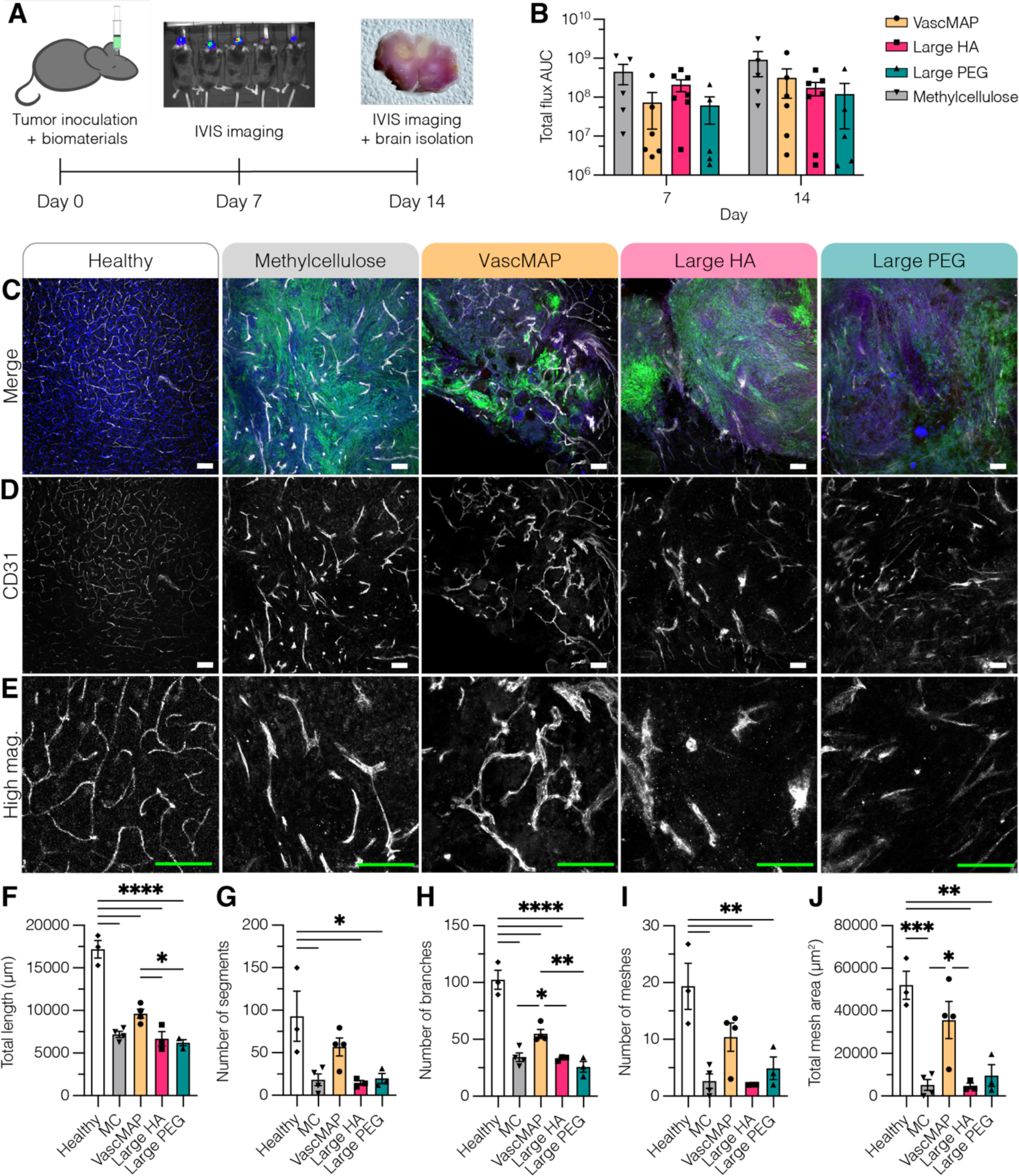
(**A**) Tumor growth was assessed at days 7 and 14 via IVIS imaging following CT2A tumors cell injections in different biomaterial matrices. Brains were isolated at Day 14 for sectioning and immunofluorescence staining. (**B**) Total flux reported as area under the curve (AUC) over a 15-minute time series study. Two-way ANOVA with Tukey HSD was performed to compare groups (*n* = 5-7). (**C**) Immunofluorescence staining of DAPI (blue), CT2A-GFP (green), and CD31 (white) in 80 µm floating sections. (**D**) CD31 channel from 80 µm floating sections. (**E**) High magnification of CD31 staining. Scale bar = 100 µm. Quantification of vessel architecture included (**F**) total length, (**G**) number of segments, (**H**) number of branches, (**I**) number of meshes, and (**J**) total mesh area. One-way ANOVA with Tukey HSD was performed to compare groups. Significance was reported at *p* < 0.05 (*), <0.01 (**), <0.005 (***), and <0.001 (****).

After 14 days of tumor growth, we isolated the brains for immunofluorescence staining to assess the vascular architecture in 80 μm floating sections for each material condition. We stained for perfused vessels via tomato-lectin (**Figure S5A-C**) as well as both perfused and non-perfused vessel structures through PECAM-1 (CD31) staining (**Figure 7C-E**). For both markers, we observed thicker vessel structures in all tumor groups compared to the healthy vascular endothelium (not quantified) (**Figure 7**, **Figure S5**). The healthy vascular endothelium boasted significantly longer perfused vascular networks with more branches compared to all tumor groups (**Figure S5D-H**). Though we observed trends of enhanced segments, meshes, and mesh area in the perfused vascular networks of VascMAP, there were no significant differences compared to the other biomaterial groups (**Figure S5D-H**).

However, when considering both perfused and non-perfused vessel structures, we observed greater differences amongst groups. In comparison to healthy brain tissue, all tumor groups had less total vessel length and number of branches (**Figure 7F-G**). However, tumors with VascMAP had greater total vessel length than the other two MAP groups, and VascMAP had more branches than all other tumor groups (**Figure 7F-G**). In terms of the number of vessel segments (either connected or disconnected), VascMAP was the only group which did not vary significantly from healthy tissue (**Figure 7H**). In the VascMAP condition, we observed vessel structures wrapping around the µgels in the tumor and forming loops. When we quantified meshes, or loops, the trends show VascMAP had more meshes than other tumor groups (though not statistically significant); however, VascMAP was the only group that did not vary significantly from healthy brain tissue for meshes and total mesh area (**Figure 7I-J**). By using this murine model of glioblastoma, we were able to show that VascMAP alters the development of tumor vasculature by instructing differential vascular patterning.

## 3. Conclusion

We modulated both the microstructure of our MAP scaffold platform and the spatial bioactivity to promote de novo assembly of endothelial progenitor-like cells into vessel-like structures. Microstructure was tuned by altering µgel size, while spatial bioactivity was controlled using heterogeneous μgel populations. Through a combination of *in silico* and *in vitro* experimentation, we found that microstructure (by way of void space porosity), integrin binding, and growth factor sequestration were all shown to guide vascular morphogenesis. We first determined that MAP scaffolds with pore volumes on the same order of magnitude as cells facilitate cell growth and vacuole formation. We then fabricated scaffolds using well-mixed combinations of heterogeneous μgel populations to control the spatial presentation of bioactive cues. Both μgel size and proportion were shown to have an effect on the spatial presentation of bioactive cues. In terms of cell response, we achieved spatial control over cell spreading by incorporating adhesive μgels in heterogeneous MAP scaffolds. While we demonstrated that integrin engagement was the primary driver of network formation in these materials, cell growth was directed to μgels loaded with heparin nanoparticles when all μgels in the system were adhesive. Heterogeneous MAP scaffolds comprising 50% RGD + 50% nH/RGD μgels with an average μgel diameter of 80 μm lead to robust cell networks after 3 days in culture with phenotypes resembling vascular structures boasting lumens and a basement membrane. We finally showed that the findings elucidated by our reductionist *in vitro* model translated to *in vivo* vessel formation. In a dermal wound healing model, the unique VascMAP scaffold formulation supported enhanced vessel formation and long-term vessel maturation. In a glioblastoma tumor model, VascMAP instructed differential vascular patterning in the tumor microenvironment. Taken together, this work determined a MAP scaffold formulation with µgels of specific size, adhesivity, and growth factor-trapping capability that led to primitive vascular constructs *in vitro* and facilitated vessel formation *in vivo*.

## 4. Experimental Methods

### Material synthesis

Hyaluronic acid (HA) (Contipro, pharmaceutical grade, 79 kDa) was modified at the carboxylic acid site with 5-norbornene-2-methylamine (NB) (TCI Chemicals) as described previously.^[29]^ Approximately 35% of HA repeat units were successfully modified with NB, as determined by proton NMR analysis performed in deuterium oxide. ^1^H NMR shifts of pendant norbornenes at δ6.33 and δ6.02 (vinyl protons, endo), and δ6.26 and δ6.23 ppm (vinyl protons, exo) where compared to the HA methyl group δ2.05 ppm to determine functionalization. HA-tetrazine was synthesized as described previously using 79 kDa HA with molar equivalents of 1:1:0.25 of HA-repeat units to 4-(4,6-dimethoxy-1,3,5-triazin-2-yl)-4-methylmorpholinium chloride (DMTMM) (Thermo Fisher Scientific) to tetrazine-amine (Chem-Impex).^[16]^ ^1^H NMR shifts of pendant tetrazine groups at δ8.5 (2H) and δ7.7 (2H) (aromatic protons) were compared to the HA methyl group δ2.05 ppm to determine functionalization. Alexa Fluor 488 C5-tetrazine (Alexa488-Tet) and Alexa Fluor 555 C2-tetrazine (Alexa555-Tet) were synthesized through two base-catalyzed thiol-Michael addition reactions in series, as described previously.^[30, 31]^ Heparin sodium salt (14,000 Da; EMD Millipore) was modified with norbornene functional handles and fabricated into nanoparticles as described previously by Wilson, et al.^[21]^ ^1^H NMR shifts of pendant norbornene groups at δ6.33 and δ6.02 (vinyl protons, endo) and δ6.26 and δ6.23 ppm (vinyl protons, exo) were compared to the heparin methyl group at δ2.00 ppm to determine functionalization.

### HA-NB Microgel Production

Microfluidic production of HA-NB μgels has been described previously.^[30]^ In brief, μgels formulated using HA-NB (3.4 wt% (w/v)) precursor solution in 0.3 M HEPES buffer with 3.5 mM MMP-cleavable crosslinker (Ac-GCRDGPQGIWGQDRCG-NH2, GenScript), 1.75 mM Tris(2-carboxyethyl)phosphine hydrochloride (TCEP, Millipore Sigma), and 9.9 mM lithium phenyl-2,4,6-trimethylbenzoylphosphinate (LAP, TCI Chemicals). For RGD-modified μgels, 1 mM RGD peptide (GenScript) was added to the precursor solution. For μgels with heparin nanoparticles, the nanoparticle stock was added for a final concentration of 0.167 μg per μL precursor solution. The surfactant/oil phase for the microfluidic μgel generation consisted of 5% (v/v) Span-80 (Millipore Sigma) in heavy white mineral oil (Avantor) degassed prior to use. The μgel precursor solution was sterile filtered using a 0.2 µm filter then loaded into a 1 mL syringe alongside a 5 mL syringe of oil/surfactant on a dual-syringe pump run at 0.3 µL min^-1^. The μgel production was conducted on a microfluidic device cast with Sylgard 184 polydimethylsiloxane (PDMS) (Dow Corning) prepared and cured according to the manufacturer’s instructions then plasma bonded to a glass slide using a corona plasma gun. Prior to the run, Rain-X was flushed through the device to coat the channels. The μgels fabricated on the device were collected in a 15 mL conical tube with a UV light fixed over the collection tube to facilitate downstream crosslinking at an intensity of 20 mW cm^-1^. The μgels were purified by first washing with sterile-filtered 2% Pluronic F-127 (Sigma-Aldrich) (w/v) in washing buffer (300 mM HEPES, 50 mM NaCl, 50 mM CaCl_2_) followed by repeated washes (centrifugation (5,000 RCF) and aspiration of the wash) with washing buffer. All washes were performed with sterile-filtered buffers in a sterile hood. The μgels were fluorescently labeled using Alexa Fluor-Tetrazine conjugates. After staining, microgels were washed 3x with 1x PBS. The purified μgels were washed with 70% ethanol to sterilize. The μgels were incubated 1:1 in 70% ethanol overnight then lyophilized.

### Simulations/LOVAMAP

Methods for simulations and LOVAMAP software follow those described in Riley, et al.^[17]^ Briefly, SideFX Houdini software (rigid-body solver) was used to simulate MAP scaffolds that contained rigid spheres. All parameters other than sphere diameter were held at default settings for all runs. Particles were randomly initialized above a 600 × 600 × 600 µm^3^ container and dropped into the container through a funnel to obtain random packing. Simulated scaffold data (particle centers and radii) were then inputted into custom LOVAMAP software for analysis of 3D pore volume, gap size, and paths. LOVAMAP uses particle configuration and Euclidean distance transforms (EDTs) of the void space to identify spatial landmarks (e.g., local center points of space) that are used to segment the space into 3D pores. For **Figure 1F**, particle diameters were sampled from distributions in **Figure 1B** (*N* = 10 simulated scaffolds per group), and pore volumes of interior pores were reported in picoliters. To assess gap size in **Figure 2K-N**, particles representing Population B were randomly removed at different proportions from a simulated scaffold comprising 60 µm diameter particles (*N* = 10 repeated measures per group), and the spanning diameter at each local center point (derived from the EDT) is reported to capture gap length between remaining Population A particles. Regarding path data in **Figure 3A-J**, LOVAMAP outputs all shortest paths from the center of the scaffold toward exit points along the medial axes. Paths travel through consecutive pores, and so-called adhesive-µgel paths are defined as paths where every pore along the path is surrounded by at least one adhesive particle. In contrast, heterogeneous-µgel paths are defined as paths where every pore along the path is surrounded by at least 40% and at most 60% of adhesive particles. Particles in a simulated scaffold were randomly chosen as adhesive particles at different proportions (*N* = 10 repeated measures per group) and proportion of adhesive- or heterogeneous-µgel paths were reported.

### hECFC culture

hECFC culture has been described previously by Kusuma, et al.^[23]^ In brief, hECFCs were cultured according to manufacturer’s instructions in endothelial growth medium 2 (EGM2, Lonza) containing 10% fetal bovine serum (FBS) on type I collagen-coated flasks (BD Biosciences).

### ECFC culture in MAP scaffolds

To create a custom cell culture device for these experiments, a negative mold was 3D printed using a 3D, Form 2 stereolithography printer (Formlabs, Inc., Duke SMIF).^[30]^ The culture wells were composed of a lower cylindrical culture section (4 mm in diameter and 1 mm tall) and a larger cylindrical media reservoir (5 mm in diameter and 3 mm tall) above the lower culturing section able to contain at least 50 µL of media. The wells were cast with PDMS and bonded to glass coverslips then autoclaved for sterilization. The lyophilized μgels were rehydrated at 3 wt% (w/v) in ECFC media. HA-Tet was dissolved at 0.02 mg mL^-1^ in EGM medium for all conditions in a final volume of 16%. Cells were seeded at 5,000 cell µL^-1^ MAP. μgels and HA-Tet were mixed well with the cell pellet and seeded at 5 µL per well. Interlinking occurred at 37°C for 25 minutes prior to the addition of excess medium, and media was changed daily. For functional antibody blocking experiments, either CD44 monoclonal antibody (IM7, eBioscience) or Integrin beta1 antibody concentrate (AIIB2, Developmental Hybridoma Bank) were diluted in the cell media to 20 μg/mL or 5 μg/mL, respectively, and 75 μL were added to each well. At time points of either 2, 24, or 72 h the wells were fixed with 4% paraformaldehyde for 30 minutes at room temperature then washed three times with 1x PBS.

### Immunofluorescent staining of ECFC cultures in MAP scaffolds

Samples were blocked with 0.15% triton-x in PBS for 1 hour at room temperature. The wells were stained with DAPI (1:1000) (Sigma-Aldrich) and Alexa Fluor™ 647 Phalloidin (1:40) (Invitrogen) in the same blocking buffer overnight at 4°C. For staining of the extracellular matrix components, samples were blocked with 5% normal goat serum and 0.15% triton-x in PBS for 1 hour at room temperature. Samples were incubated with primary antibodies for collagen IV (Abcam), fibronectin (Sigma Aldrich), or laminin (Abcam) 1:100 for 1 hour at room temperature. The samples were washed three times with 1x PBS then incubated in secondary antibodies (Invitrogen) 1:500 for 2 hours at room temperature. Samples were then washed three times with PBS and imaged on a confocal microscope at 20x and 40x magnification (Z-stacks with 2.5 um step size).

### Image analysis of MAP scaffolds

In IMARIS software, the spot counter tool was used to quantify the number of μgels as well as identify their X, Y, and Z coordinates for each population using the corresponding fluorescent signal. A custom MATLAB script was written to calculate the distance between μgels using the X, Y, and Z coordinates and determine the distance to the nearest μgel. Another custom MATLAB script was written to analyze the fluorescence of each μgel population within 150 μm x 150 μm x 150 μm volumes throughout the Z-stack.

### Image analysis of cell response

In IMARIS software, 3D volume renderings of confocal Z-stacks were used to quantify cell volume (F-actin) and scaffold volume. The IMARIS spot counter tool was used to quantify cell counts using DAPI signal. Number of touching cells and number of vacuoles were quantified manually in ImageJ. The ImageJ plugin DiaAna was used to quantify colocalization of nuclei (DAPI signal) with fluorescently labeled µgel populations. The ImageJ Angiogenesis Analyzer plugin was used to assess ECFC network length and branches. ImageJ was used to quantify % area of the fluorescent channel for each extracellular matrix component.

### PEG MAP scaffold fabrication

The fabrication of the PEG μgels used in this work has been described previously by Liu, et al. ^[32]^ The precursor solution for these μgels included 8-arm PEG Vinylsulfone (PEG-VS), K-peptide (Ac-FKGGERCG-NH2, GenScript), Q-peptide (Ac-NQEQVSPLGGERCG-NH2, GenScript), and RGD (Ac-RGDSPGERCG-NH2, GenScript) in 0.3 m triethylamine (Sigma) buffer. The crosslinker solution was prepared by dissolving the MMP-cleavable peptide (Ac-GCRDGPQGIWGQDRCG-NH2, GenScript) in distilled water at 12 mm and 10 μm Alexa-Fluor 647-maleimide (Invitrogen). These solutions were filtered through a 0.2 μm sterile filter before loading into 1 mL syringes. The final composition of μgels were 5% (w/v) PEG-VS with 500 um K-peptide, Q-peptide, and RGD, respectively. For large μgel generation, a dual-inlet device with one for the aqueous solution (pre-mixed the precursor solution with an equal volume of the crosslinker solution) and the oil phase (heavy mineral oil with 5% v/v Span-80), was used. The μgels were collected in a Span-80 (5% v/v) and triethylamine (3% v/v) oil bath and allowed to gel overnight at room temperature. These μgels were then purified by repeated washes with a HEPES buffer (0.3 m, pH 8.3 containing 1% Antibiotic– Antimycotic and 2% Pluronic) and centrifugation. To form MAP scaffolds with PEG-VS μgels, 1 μL thrombin (200 U per mL in 200 mm Tris-HCl, 150 mm NaCl, 20 mm CaCl2) and 2 μL Factor XIII (250 U per mL) were combined with 50 μL of μgels, mixed via thorough pipetting, and allowed to incubate at 37 °C for 30 min to form a solid hydrogel.

### Wound healing model

The assessment of biomaterials in dermal wounds was performed using a standardized wound healing model that has been described previously.^[33]^ In brief, both male and female SKH-1 mice (age 10-16 weeks) received four 5-mm full thickness splinted wounds on their back followed by 10 µL injection of each biomaterial. Wounds were macroscopically imaged over two weeks to assess the wound closure rate. Each wound was excised using an 8 mm biopsy punch around the original wounding area and then cut in half, with one side embedded in paraffin and the other in optimal cutting temperature (OCT) compound.

### Wound immunofluorescence

Tissue samples embedded in OCT were cut into 20 µm sections on a cryostat. Skin sections were fixed with acetone for 10 min then washed three times with 1×PBS + 100 mM CaCl2 + 100 mM MgCl for 10 min at room temperature. For CD31 and NG2 staining, slides were incubated in a blocking solution containing 0.3% Triton X-100 and 10% normal donkey serum (EMD Millipore) in 1xPBS + 100 mM CaCl2 + 100 mM MgCl for 1 h at room temperature. For Keratin-14 staining, separate slides were incubated in a blocking solution containing 0.3% Triton X-100 and 10% normal goat serum (Abcam) in 1xPBS + 100 mM CaCl2 + 100 mM MgCl. The slides were then incubated in the primary antibodies (rat anti-PECAM-1, clone 390, EMD Millipore; rabbit anti-NG-2 chondroitin sulfate, EMD Millipore; rabbit anti-Keratin-14, BioLegend) diluted 1:100 in the blocking solution overnight at 4 °C. The slides were washed three times with 1×PBS + 100 mM CaCl2 + 100 mM MgCl for 10 min at room temperature. The slides were then incubated in the secondary antibodies (ThermoFisher) and nuclear marker DAPI (Sigma-Aldrich) diluted 1:500 in the blocking solution for 2 h at room temperature. The slides were washed three times with 1×PBS + 100 mM CaCl2 + 100 mM MgCl for 10 min then allowed to dry at room temperature. The slides were dehydrated in ascending concentrations of ethanol (50-100%), incubated in xylene, and mounted in mounting medium (DPX, Sigma-Aldrich). A Nikon-C2 laser scanning confocal microscope with 4x, 10x, and 20x air objectives was used to take fluorescent images represented as maximum intensity projections. CD31 expression was quantified in the wound area using ImageJ. The colocalization of CD31 and NG-2 fluorescence was quantified using Just Another Colocalization Plugin (JaCoP) in ImageJ.^[34]^ Epidermis thickness was quantified by manually tracing the region positive for Keratin-14 in the wound then dividing by the length. All measurements were averaged across 2-3 sections from each wound and are presented per animal.

### Wound immunohistochemistry

Tissue samples were fixed for 4 hours at room temperature in 4% PFA before paraffin embedding. 5 µm thick sections were then cut using a microtome (Epredia). Slides were de-waxed and re-hydrated using xylene and decreasing ethanol concentrations then blocked with peroxidase block (OriGene) for 10 minutes at room temperature. Sections were washed 3X in deionized water before performing Heat Induced Epitope Retrieval in sodium citrate buffer (10 mM Sodium citrate, 0.05% Tween 20, pH 6.0) at 95°C for 20 minutes. Slides were cooled for 10 minutes under running tap water and washed 3X in PBS-T. Sections were blocked with protein block (OriGene) for 10 minutes at room temperature. The primary antibody incubation at 4°C overnight immediately followed this. To stain for vasculature, sections were incubated in Armenian Hamster anti-CD31 (Developmental Studies Hybridoma Bank, 2H8). To stain for fibrosis via myofibroblast presence, sections were incubated in Rabbit anti-Alpha Smooth Muscle Actin (ABCAM, ab5694). Sections were then developed using the respective Polink-2 Plus HRP with DAB kit (Origene, D87-6 Hamster D39-6 Rabbit). Sections were counterstained in Hematoxylin (EMS) and then cleared and dehydrated using increasing ethanol concentrations followed by xylene. Sections were mounted in DPX mounting media (EMS) and then imaged at 20X using an AxioScan.Z1 (Zeiss). Images were analyzed using HALO (Indica Labs). The Indica Labs - Area Quantification v2.4.3 plugin was used to identify DAB positive areas for each wound region to give %DAB positive.

### Tumor inoculations with biomaterials

CT2A murine glioma cells expressing GFP and firefly luciferase were graciously donated by Dr. Yvonne Chen’s lab at UCLA. The CT2A cells were cultured in Dulbecco’s Modified Eagle Medium (DMEM) media supplemented with 10% fetal bovine serum and 1% penicillin/streptomycin and expanded for 3 days in culture prior to tumor inoculation. Our murine model of GBM was adapted from Swan, et al. with modified injection coordinates and a homogeneous tumor composition with lowered cell density.^[35]^ 6- to 12-week-old female C57B6/J mice (Jackson labs) were used in accordance with the approved Duke Institutional Animal Care and Use Committee (IACUC) protocol. Mice were anesthetized using 3 to 5% isoflurane given subcutaneous injection of Buprenorphine SR at 1 mg/kg. The head of the mouse was shaved then sterilized. A small incision was made followed by adding 1-2 drops of 0.25% bupivacaine locally. A stereotactic device was used to center the Hamilton syringe at the crosshairs of bregma. We injected the cell/biomaterial mixture at 0.5 mm posterior and 1.5 mm left of bregma and 2.5 mm deep. A Hamilton syringe with a 25-gauge needle connected to a syringe pump was used to inject 5 µL each biomaterial condition with tumor cells at 1 µL min^-1^. The syringe was removed incrementally over 3 minutes to prevent the material from coming out of the injection site. The site of injection was sealed with bone wax (Lukens), and the incision was closed with Vetbond (3M).

Following the surgical procedure, animals were given supplemental food and monitored every two days for signs of distress or illness. At 7- and 14-days following tumor injection, animals were imaged with IVIS Kinetic (Caliper Life Sciences) using an intraperitoneal injection of 0.2 μm filtered Xenolight D-luciferin (Perkin Elmer) at 150 mg/kg body weight. 12 minutes post-injection animals were imaged with a time series study in 5-minute increments over 15 minutes.

### Immunofluorescent staining of floating sections

After IVIS imaging on day 14, animals were euthanized and perfused with 1xPBS followed by 4% paraformaldehyde (PFA). The brains were isolated and incubated in 4% PFA overnight at 4°C. Brains were then transferred to 30% sucrose solution with 0.1% proclin and kept at 4°C until the brains sank to the bottom of the container. The brains were then sectioned on a cryostat at a thickness of 80 µm then transferred to a 24-well plate with 1xPBS.

Immunofluorescent staining was conducted in the 24-well plate using a working volume of 250 µL. Samples were blocked with 10% donkey serum in 0.3% Triton-X in PBS overnight at 4°C. For CD31 staining, samples were incubated 1:100 with anti-CD31 (rat host; Santa Cruz) in the blocking solution and incubated for 7 hours at room temperature. Samples were washed 3 times with PBS prior to overnight incubation at room temperature with secondary antibody and DAPI. For tomato lectin staining, samples were incubated 1:500 with Streptavidin Alexa Fluor™ 647 Conjugate (Vector Laboratories) overnight at 4°C in 1xPBS. All samples were washed in PBS then transferred to glass slides for imaging on a confocal microscope (Nikon Ti Eclipse).

### Statistical analysis

For *in vitro* and *in silico* experiments, data are presented with error bars representing standard deviation. For *in vitro* experiments, *n* = 3 biological replicates with each performed with duplicate technical replicates. Two-way ANOVA was performed on *in vitro* experiments to compare across multiple time points. Upon significance of the interaction term, Tukey HSD post-hoc analysis was performed to compare conditions. For the following experiments, one-way ANOVA was performed which yielded *P* < 0.05 and prompted post-hoc analysis (Tukey HSD): distance to nearest µgel, vacuole formation after 24 hours, and the comparison of conditions at 72 hours for nuclei, total volume, network length, and branches. For *in vivo* experiments, data are presented with error bars representing standard error. For the wound healing experiments, *n* = 6 animals with 3 female and 3 male mice for each time point. Animals were only excluded from analyses if the wound could not be located in the skin sections. For the tumor experiments, the number of animals analyzed for each group varied depending on survival of the inoculation procedure. Two-way ANOVA was performed to compare groups across time points. Upon significance of the interaction term, Tukey HSD post-hoc analysis was performed to compare conditions. For all experiments, significance is indicated by \**P* < 0.05, \*\**P* < 0.01, \*\*\**P* < 0.001, \*\*\*\**P* < 0.0001. Statistical analyses were performed with GraphPad Prism software.

## Acknowledgements

We would like to thank Dr. Katrina L. Wilson and Kevin Erning for the synthesis of heparin nanoparticles as well as Dr. Yining Liu for the synthesis of the PEG microgels used in this work. We would like to thank Dr. Peter Cheng for simulating the particle domains analyzed with LOVAMAP. We would like the thank Dr. Sheridan Swan for her time teaching the glioblastoma tumor model that was adapted for this work. We would like to thank Dr. Yvonne Chen’s lab at UCLA for providing the glioma cells for this work. Lastly, we would like to thank Dianne Cruz and the White-McGarrah Lab at Duke University for their generosity in allowing us to use their AxioScan.

