## Supporting Information for "Engineering the microstructure and spatial bioactivity of MAP scaffolds in vitro instructs neovascularization in vivo"

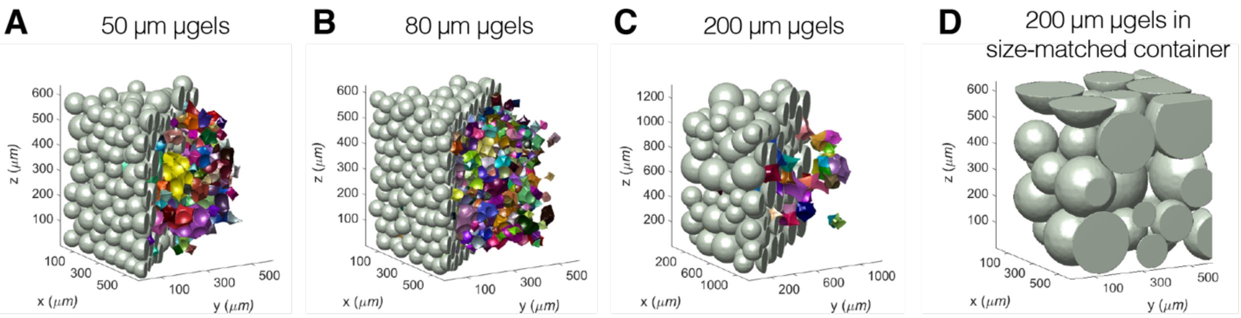


**Figure S1.** Simulated scaffolds of rigid spheres (gray) to represent MAP scaffolds with average diameters of (**A**) 50 µm, (**B**) 80 µm, and (**C**) 200 µm µgels shown as half of the scaffold with the pores determined by LOVAMAP shown in color. (**D**) Simulated scaffolds of rigid spheres to represent MAP scaffolds with 200 µm µgels in a size-matched container to the simulated scaffolds in (**A**) and (**B**).

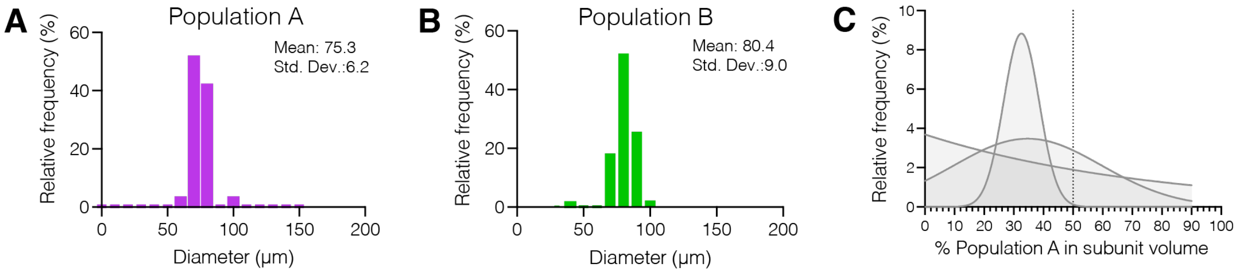


**Figure S2.** (**A-B**) Frequency distributions of the µgel diameters used in heterogeneous MAP scaffold experiments. (**C**) Frequency distribution of the percentage of Population A within a subunit volume of the confocal Z-stack of three independent MAP scaffolds composed of 50% A + 50% B with poor mixing.

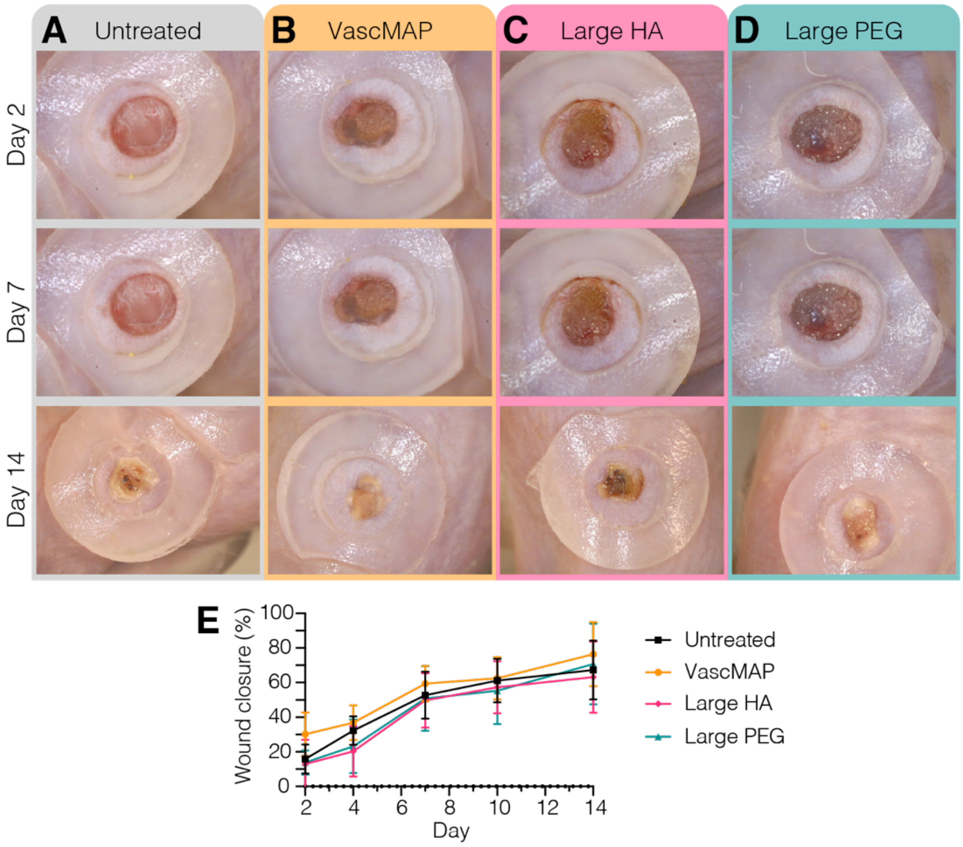


**Figure S3.** Wound closure for the dermal wound healing model was assessed over 14 days. Representative macroscopic images of each wound from the same mouse are shown for (**A**) untreated, (**B**) VascMAP, (**C**) Large HA MAP, and (**D**) Large PEG MAP at days 2, 7, and 14. (E) Quantification of wound closure over 14 days. Repeated measures ANOVA did not yield significance.

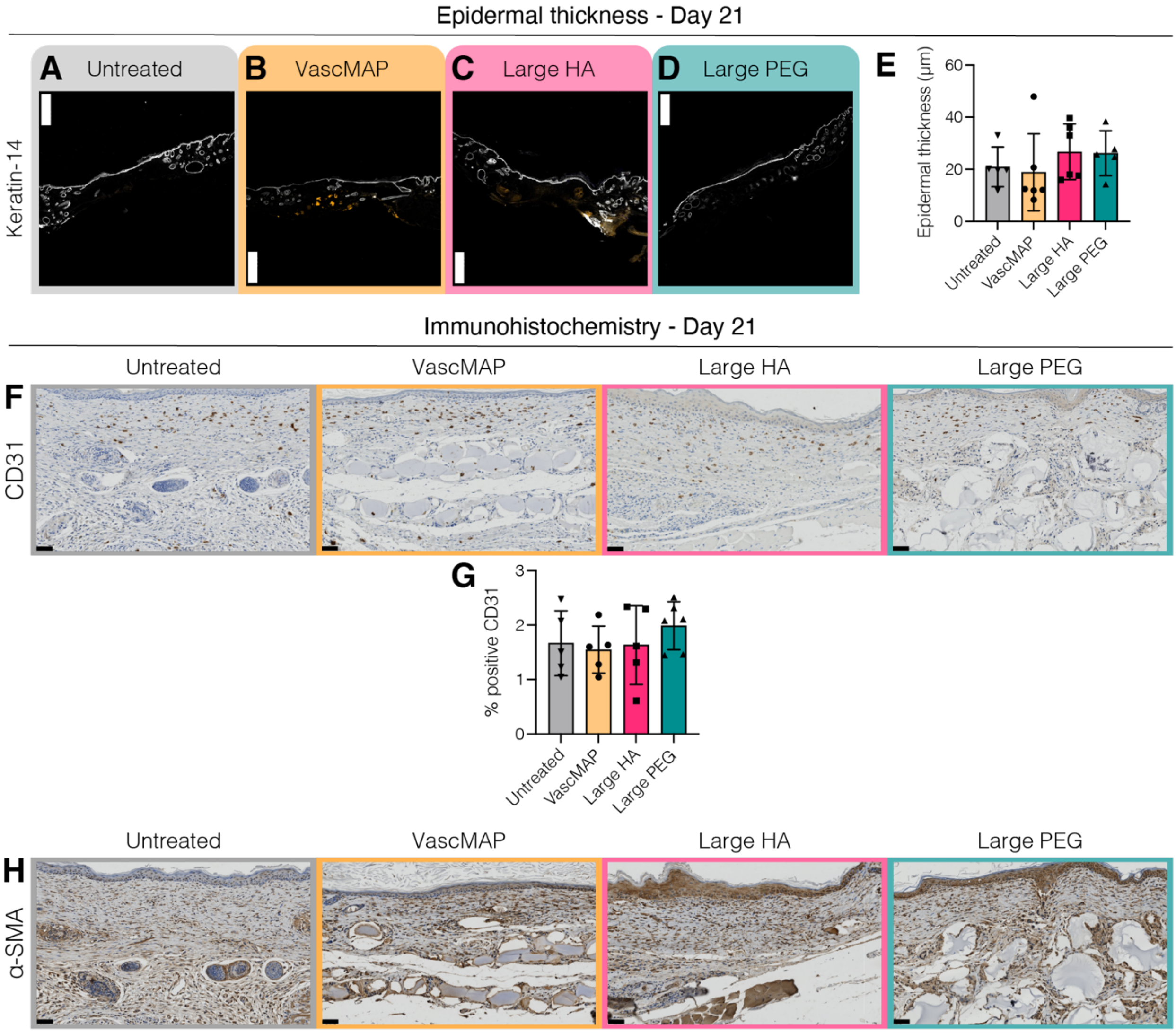


**Figure S4.** Epidermal thickness was quantified at Day 21 using positive keratin-14 staining in the wound region for (**A**) untreated, (**B**) VascMAP, (**C**) Large HA MAP, and (**D**) Large PEG MAP. (**E**) Quantification of epidermis thickness. One-way ANOVA with Tukey HSD was performed to compare groups. (**F**) In addition to immunofluorescence staining, immunohistochemical (IHC) staining for CD31 was also performed on Day 21. (**G**) Quantification of CD31 from IHC reported as % positive area of CD31. One-way ANOVA with Tukey HSD was performed to compare groups. Significance was reported at $p < 0.05$ (*), <0.01 (**), <0.005 (***), and <0.001 (****). (**H**) IHC staining for α-SMA on Day 21.

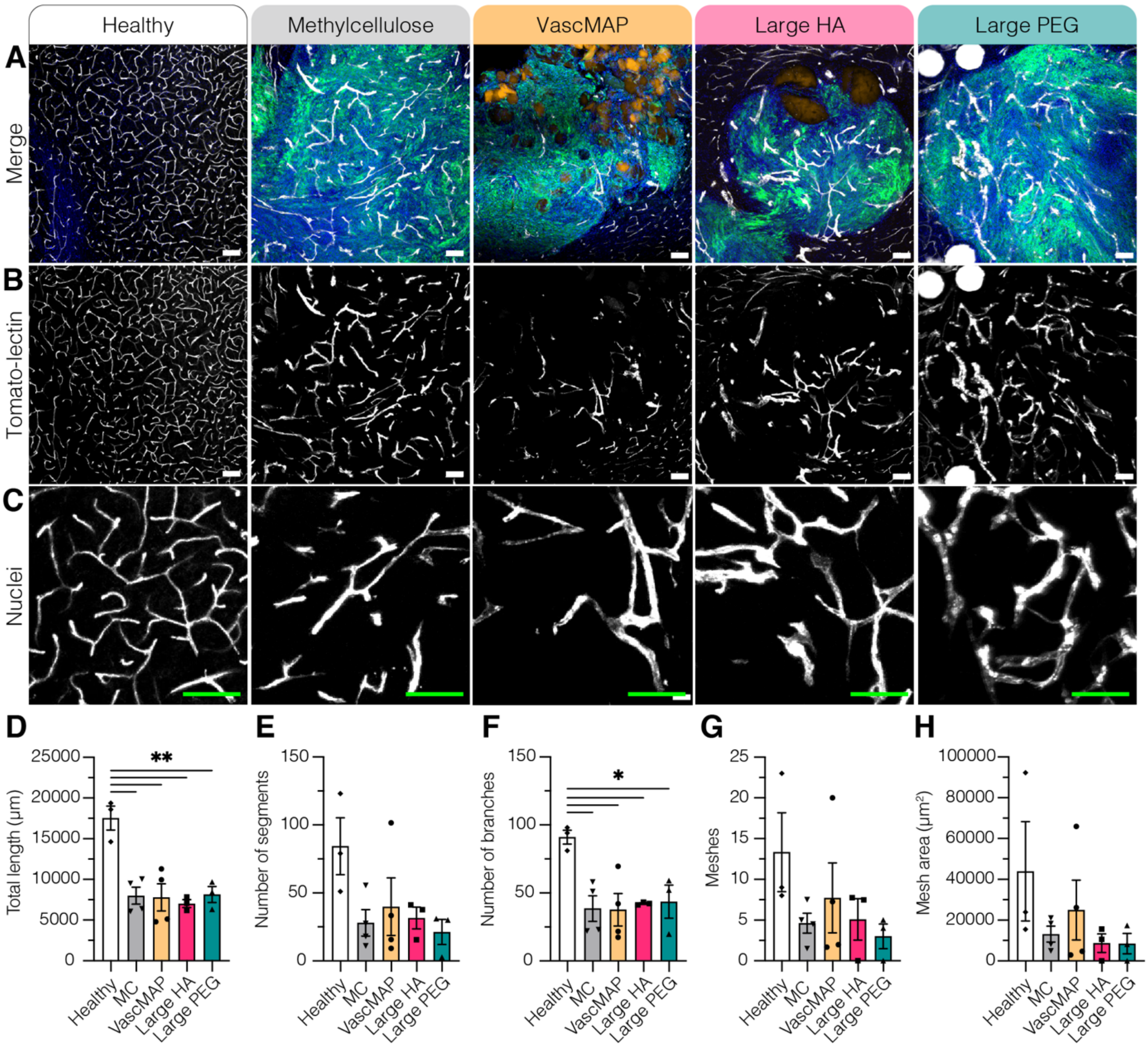


**Figure S5.** (**A**) Immunofluorescence staining of DAPI (blue), CT2A-GFP (green), MAP scaffolds (green/orange), and tomato-lectin (white) in 80 µm floating sections. (**B**) Tomato-lectin channel from 80 µm floating sections at 20x magnification. Scale bar = 100 µm. (**E**) High magnification of tomato-lectin staining. Scale bar = 100 µm. Quantification of vessel architecture included (**F**) total length, (**G**) number of segments, (**H**) number of branches, (**I**) number of meshes, and (**J**) total mesh area. One-way ANOVA with Tukey HSD was performed to compare groups. Significance was reported at $p < 0.05$ (*), <0.01 (**), <0.005 (***), and <0.001 (****).
